# PROTGOAT : Improved automated protein function predictions using Protein Language Models

**DOI:** 10.1101/2024.04.01.587572

**Authors:** Zong Ming Chua, Adarsh Rajesh, Sanju Sinha, Peter D. Adams

## Abstract

Accurate prediction of protein function is crucial for understanding biological processes and various disease mechanisms. Current methods for protein function prediction relies primarily on sequence similarities and often misses out on important aspects of protein function. New developments in protein function prediction methods have recently shown exciting progress via the use of large transformer-based Protein Language Models (PLMs) that allow for the capture of nuanced relationships between amino acids in protein sequences which are crucial for understanding their function. This has enabled an unprecedented level of accuracy in predicting the functions of previously little understood proteins. We here developed an ensemble method called PROTGOAT based on embeddings extracted from multiple and diverse pre-trained PLMs and existing text information about the protein in published literature. PROTGOAT outperforms most current state-of-the-art methods, ranking fourth in the Critical Assessment of Functional Annotation (CAFA 5), a global competition benchmarking such developments among 1600 methods tested. The high performance of our method demonstrates how protein function prediction can be improved through the use of an ensemble of diverse PLMs. PROTGOAT is publicly available for academic use and can be accessed here: https://github.com/zongmingchua/cafa5

## Background

Proteins are molecules formed by long chains of amino acids which assume complex three-dimensional conformations and are essential to all biological activities. With the increasing pace of developments in sequencing technologies ever more protein coding sequences have been sequenced and deposited in public repositories. However, most of these protein sequences are poorly annotated and require substantial time and resources from laboratory experiments to be fully characterized. Given the gap in knowledge between the number of sequenced proteins and the known functions of those proteins, substantial efforts have been made to create tools that can automatically annotate protein functions using Gene Ontology (GO) terms to evaluate their predictions.

GO terms were developed over 20 years ago in order to describe the varied functions of genes and their protein products^1^. GO terms can be divided into three distinct GO domains : Biological Process (BP), which is the broad biological process a protein is associated with, Cellular Component (CC), which is the location in a cell where a protein carries out its function, and Molecular Function (MF), which represents the specific molecular activities a protein is associated with. GO terms are all associated with one another within the form of directed acyclic graphs (DAGs) where the nodes in DAGs are functional descriptors belonging to one of the three GO domains which are connected by relational terms.

Over the last decade or so, multiple techniques have been developed to produce automated function prediction (AFP) of proteins based on the protein primary sequence using machine learning and statistical methods^2–10^. In order to benchmark these methods against one another as well as to spur further innovation in developing new methods for predicting protein function, the Critical Assessment of Protein Function Annotation (CAFA) is regularly held with the fifth iteration (CAFA 5) having been recently concluded on Kaggle^11^. During a CAFA competition submitted models are evaluated based upon their ability to predict GO terms in one of the three GO domains for proteins that previously lacked annotations in that domain and which accumulate experimentally-validated annotations during the curation phase of the competition.

Transformer-based large language models trained on enormous amounts of text have recently shown remarkable advances in generating realistic and cogent outputs on a large variety of subject matter^12–16^. Transformer-based models have been successful in modeling sequential information like text due to their ability to learn contextual, nuanced, and long-range relationship between words. Based on the success of these large language models, efforts have been made to develop similar language models trained on protein sequences^17^. Among the best models that have emerged are the various sized ESM2 models produced by Meta and Prot-T5 by the Rost lab that have been trained on billions of protein sequences^18–20^. It has been shown that use of these language models can be an effective way to generate predictions about protein characteristics, including localization, protein-protein interactions, and function^8,21–23^. We present here PROTGOAT (PROTein Gene Ontology Annotation Tool) that integrates the output of multiple diverse PLMs with literature and taxonomy information about a protein to predict its function. PROTGOAT was ranked 4th place by maximum F1-score (F_max_) among 1600 tested methods in the latest CAFA 5 competition.

Finally, we sought to validate the predictions made by PROTGOAT through comparing the predictions made by PROTGOAT against RNA-seq data comparing proliferating and senescent cells. Cellular senescence is a conserved cellular response to stress that leads to permanent halt to replication and is an important hallmark of biological aging^24^. Distinct change in the transcriptome such as the production of a conserved, pro-inflammatory range of cytokines, chemokines, growth factors, and proteases termed the senescence-associated secretory phenotype (SASP) in senescent human cell lines has been studied extensively but remains poorly annotated in GO databases.

## Materials and methods

### Dataset

We used protein GO annotations (Swiss-Prot) from the then latest version of UniProtKB that was available from https://www.uniprot.org/help/downloads on 3 May 2023. We narrowed down the GO terms associated with each protein in our training dataset by selecting only those that were derived from experimental results (evidence code ECO:0000269, ECO:0000314, ECO:0000353, ECO:0000315, ECO:0000316, ECO:0000270), high throughput assays (evidence code ECO:0006056, ECO:0007005, ECO:0007001, ECO:0007003, ECO:0007007), annotated with traceable author statement (evidence code ECO:0000303), or inferred by curator (evidence code ECO:0000305).

Many GO terms within the training set are labeled on proteins without the necessary parental GO terms being annotated for the same protein. In our training set we added annotations for every GO label up to the root for each domain. Relationships between GO terms were obtained from https://purl.obolibrary.org/obo/go/go-basic.obo, with parental GO terms defined as GO terms that encompass

### PROTGOAT

Our model utilizes both protein sequence information as well as literature and taxonomic information to carry out novel annotations of proteins, which we have termed PROTGOAT. An overview of PROTGOAT is shown in Figure 1.

**Figure 1.**
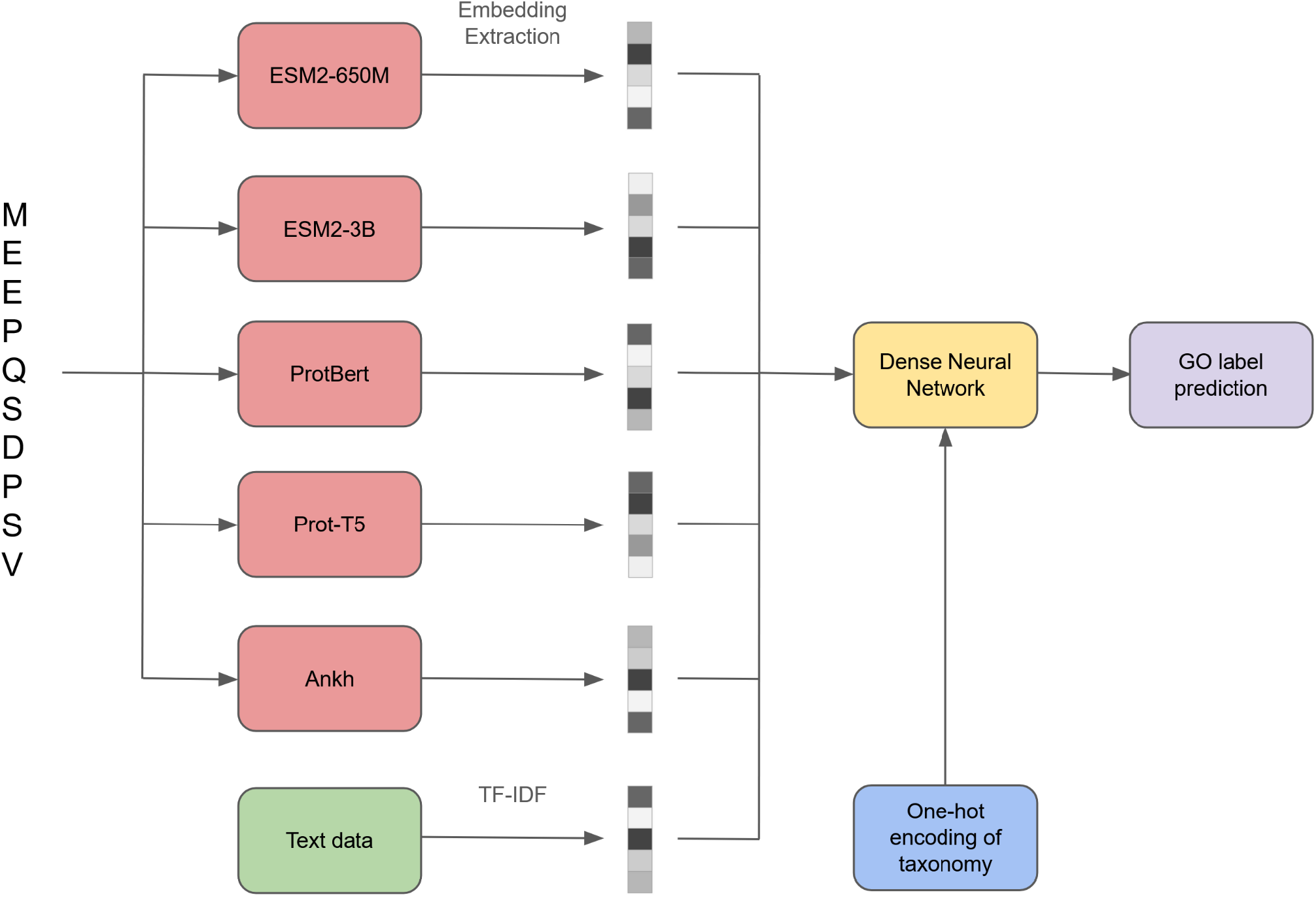
An overview of PROTGOAT, showing how multiple PLMs are used to extract different embeddings from the same protein amino acid sequence. Text data from literature that relates to the same protein together with a one-hot encoding of the taxonomy of the organism from which the protein is derived serve as separate inputs to the neural network. The final layer of the network predicts multiple GO labels for each protein while assigning a probability for each protein-GO label pair.

### Protein Language Models

We either generated or obtained from public repositories embeddings of of size 2560, 1280, 1024, 1024, and 1536 for all 142246 proteins in the training dataset using the PLMs ESM2-3B, ESM2-650M, Prot-T5, ProtBert, and Ankh respectively. Each protein in the training dataset produces an embedding of size LxN where L represents the length of the protein and N the length of the embedding representation that is unique to each PLM. We averaged values at each amino acid position of a protein to obtain a compressed 1xN size embedding representation of each protein which is used directly as the input for training our model.

### Word embedding input

We incorporate literature information into PROTGOAT as a separate input that the neural network trains on simultaneously along with embeddings derived from PLMs. A Python script was utilized to retrieve PubMed IDs for proteins in both the training and test dataset by interfacing with the UniProt REST API. This process involved querying the UniProt Knowledgebase to establish links between each protein and its related scientific literature. Following this, titles and abstracts corresponding to these PubMed IDs were extracted from the PubMed database using the Biopython Entrez utilities.

The collected abstracts were refined through a preprocessing routine using NLTK in Python. This involved removing stopwords, common words, punctuation, and numerical terms. A word count analysis was conducted to identify and exclude words appearing in more than 75% of the abstracts. This step ensured the retention of only the most relevant and informative words in the dataset, making it suitable for effective TF-IDF vectorization.

For the TF-IDF vectorization process, a comprehensive vocabulary was constructed from all Gene Ontology (GO) categories. This was achieved by using the OBO file containing GO terms and extracting all term names, which were then combined, cleaned, and tokenized. The cleaning process involved removing non-alphanumeric characters, excess whitespace, stopwords, punctuation, and numerical terms, with a focus on words longer than two characters. This led to a unique, non-redundant vocabulary representative of the entire GO term set.

To ensure consistency in the feature space, the preprocessed abstracts from both the training and test datasets were merged, with duplicates removed based on Protein ID. A TfidfVectorizer, initialized with this GO-derived vocabulary, was then used to transform the merged abstracts into TF-IDF vectors. These vectors quantified the importance of each term in the abstracts relative to the combined corpus, ensuring that both training and test datasets shared the same feature space for subsequent machine learning models. The TF-IDF vectors for each protein were then merged for the full list of train and test protein IDs, filling proteins with no text data with zeros, and then structured into a final numpy embedding for use in the final model. The final word embedding was 10279 in size.

### Training

We chose to train three distinct models, one for each of the three GO domains. Due to computational constraints, we only trained our models and made predictions for 1500 GO terms in the BPO GO domain, as well as 800 GO terms in the CCO and MFO GO domains each. GO terms were selected based upon the frequency with which the GO term appears in the training data as well as information accretion (IA) value for that GO term as defined by the competition host of CAFA 5^25^.

Many of the proteins in our training dataset lack GO terms in one or more of the three GO domains and those were excluded from the training data for the respective models of that domain. The final number of proteins in each training dataset for each individual GO domain was 93617 for BPO, 93364 for CCO, and 79132 for MFO.

We carried out training using Dense Neural Networks (DNNs) that used the embedding representation of each protein to attempt to predict GO terms for that protein. We added as additional inputs information on the taxonomy of the species from which the protein is derived using on hot encoding as well as the previously described word embeddings associated with each protein. The overall DNN architecture takes these separate inputs before a final layer that concatenates the intermediate layers of the DNN in order to produce a set of predicted GO terms for the respective protein.

We used the GO annotations for each protein as labels and binary cross entropy (BCE) as the loss function during training in order to update the model based on differences between predicted and true values, treating this as a multi-label classification problem. We employed the Adam optimizer due to its robustness, and used a batch size of 256^26^.

During training proteins are randomly split into a training dataset (80%) and a validation dataset (20%). We carried out 5-fold cross validation on the training data for all three domains to ensure that the entirety of the training data is utilized in our models. After cross validation, the probability assigned to each protein-GO term prediction is averaged across the results of all models. The averaged probability assigned to each protein-GO term prediction based on BCE loss is used for functional validation subsequently in the context of cellular senescence.

### CAFA 5 predictions

All participants in CAFA 5 are asked to generate GO annotation predictions for proteins within a test superset consisting of 141864 proteins. A smaller subset of the test superset, the test set, consists of proteins by which models are evaluated against. The test set consists of proteins that had new experimentally verified GO terms annotated during the curation phase of the competition in GO domains that those proteins previously had no annotations in.

The assigned probability of predicted terms for proteins in the CAFA 5 test superset for models generated in each domain are averaged prior to further ensembling. Finally we took the top 5% of predicted GO terms for all proteins from each domain based upon the probability assigned by PROTGOAT. The total number of GO terms predicted by PROTGOAT in the test superset were 1.1^*^10^7^, 5.7^*^10^6^, and 5.7^*^10^6^ for BPO, CCO, and MFO respectively.

## Results

### CAFA 5 results

PROTGOAT was ranked 4th on the overall leaderboard of the CAFA 5 competition as shown in Figure 2. Models were ranked based on their maximum F-measure scores by calculating weighted precision and recall for each of the GO domains in the test set of proteins and taking the arithmetic mean (BPO, CCO, and MFO)^27^. In order to capture the differences in information content between different GO terms that arise from the hierarchical nature of GO annotations, an information content (IC) value is calculated for each GO label. IC for each GO annotation is calculated by taking the logarithm of the ratio of the number of proteins with the parental terms in the ontology compared to the number of proteins with the term itself in a large pool of proteins^25^.

**Figure 2.**
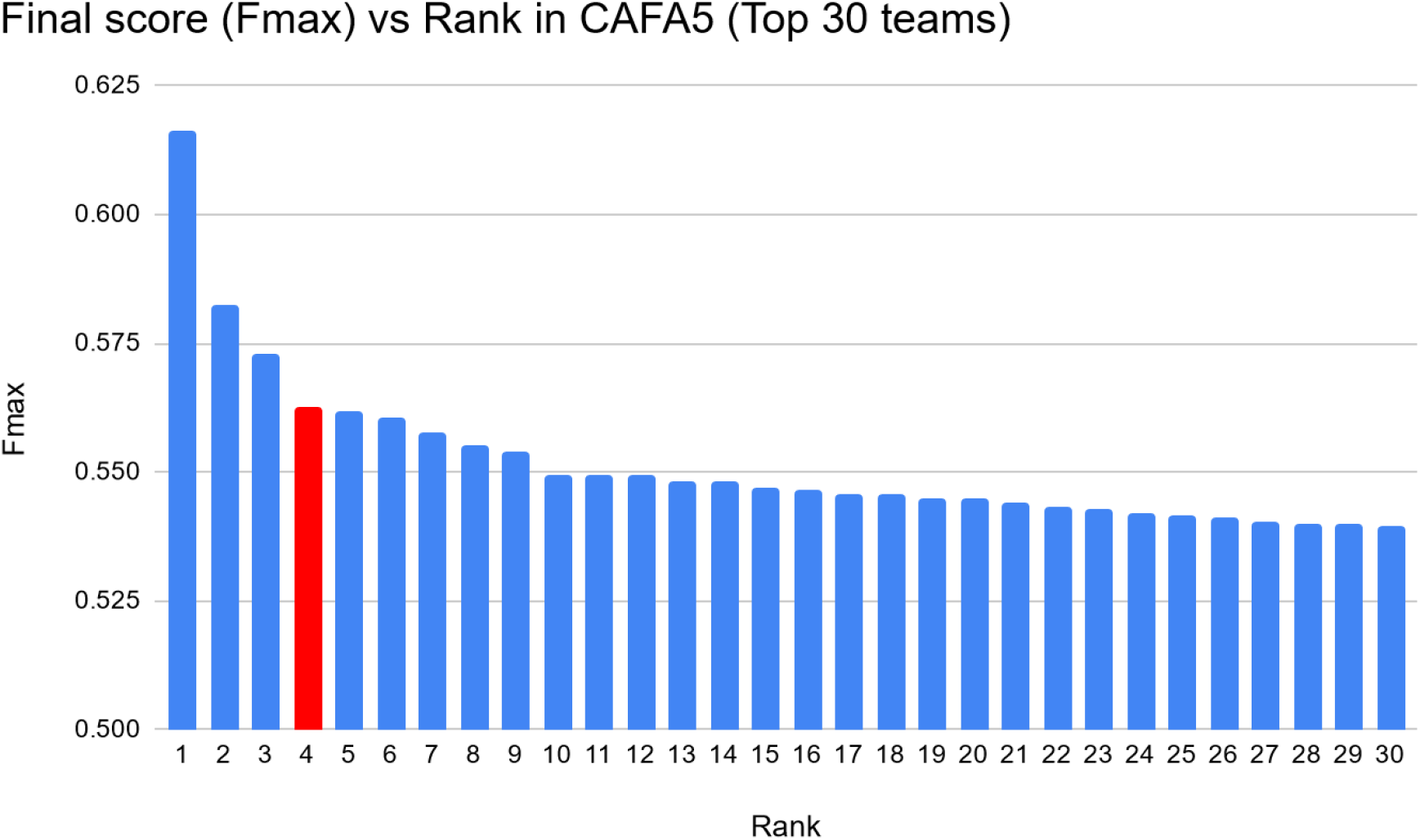
Final F_max_ scores of submitted models in CAFA 5, with PROTGOAT highlighted in red.

**Figure 3.**
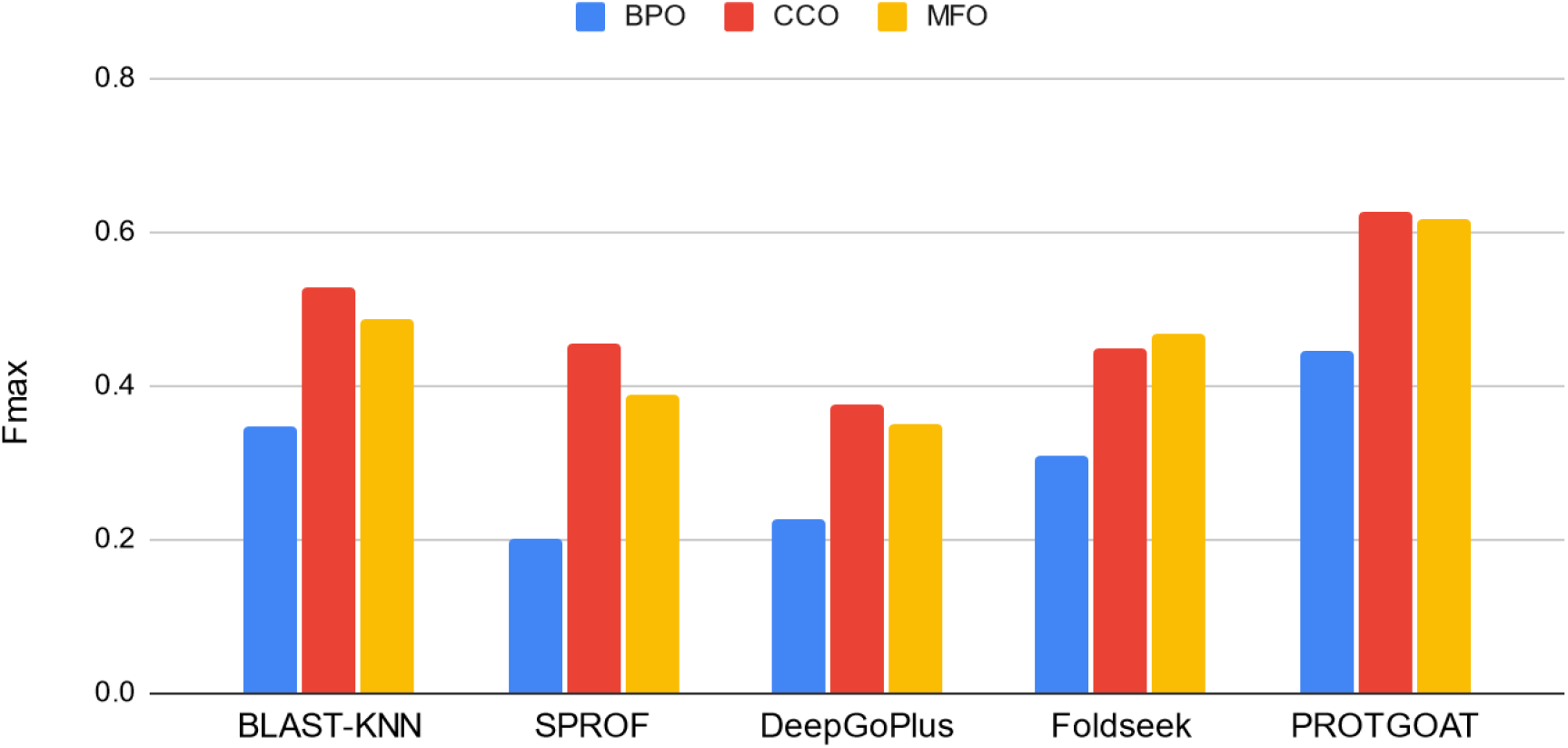
Comparison of PROTGOAT to various state of the art machine learning models for automated function prediction based on F_max_ score for each GO domain based on the final test set of CAFA 5.

### Comparison to state-of-the-art models

We compared the performance of PROTGOAT to several other state of the art machine learning models that have been released in the last few years : SPROF-GO, DeepGoPlus, and Foldseek^28–31^. We also included a BLAST-KNN model that carries out annotation transfer based on K nearest neighbor (KNN) distance based on local alignment similarity as a naive approach. We evaluated all models based on the test set of CAFA 5 and observed that PROTGOAT significantly outperforms all other models across all three GO domains.

### Ablation study

PROTGOAT utilizes various language models as well as literature and taxonomic information as training data. We carried out an ablation study to better understand the importance of each individual input for training our model. We observed that overall model performance does decline as more training data is removed. Surprisingly, we also observed that this trend is not universal for all GO domains. Predictions made for BPO GO terms declines the most steeply with decreased training data, while CCO declines less significantly. Counterintuitively, predictions made for MFO GO terms actually improves with ablation of training data, indicating that the model may be overfitting during training for predicting MFO GO annotations.

### Functional Validation

To showcase the functional efficacy of PROATGOAT, we focused on the biological process of cellular senescence (GO:0090398) as a case study. Based on the latest annotations available for genes associated with cellular senescence, there are 57 genes that have current annotations for cellular senescence and associated child terms (GO:2000772, GO:2000773, GO:2000774, GO:0090400, GO:0090402) and which are are expressed.

PROATGOAT predicted 286 human protein coding genes associated with cellular senescence process within the test superset of CAFA 5 (BCE probability > 0.05, Methods). Our predictions are enriched with annotated senescence associated genes (Fold enrichment=44.1, p-value=3.39e-51), where 35 genes are shared between predicted and known (Figure 4A).

**Figure 4.**
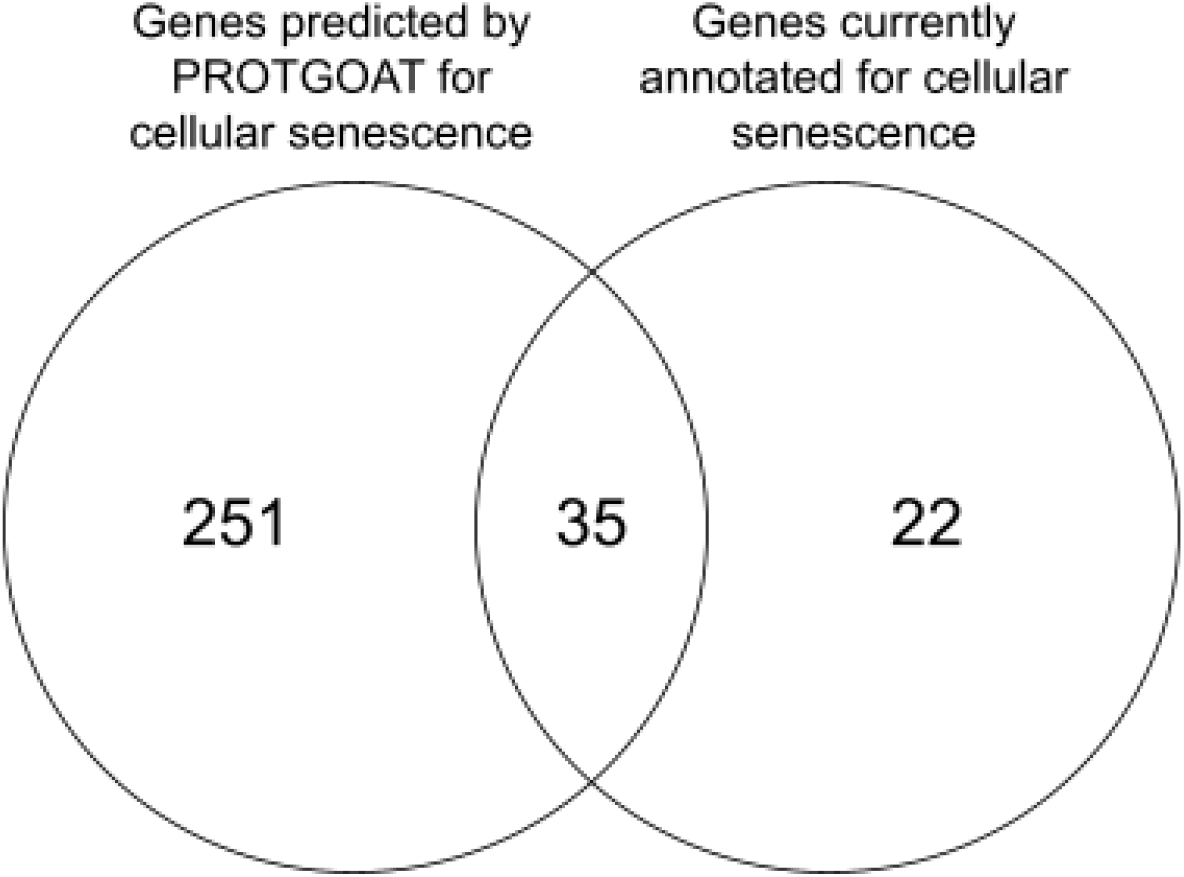
Venn diagram of number of genes associated with cellular senescence that overlap between those predicted by PROTGOAT and those that are currently annotated.

We next carried out bulk RNA-seq of IMR90 human primary fibroblasts under proliferating and senescent conditions and compared their gene expression patterns. We observed that of the 23516 genes expressed, 7190 are differentially expressed (DE) in senescent cells compared to proliferating ones (FDR P<0.01). We observed that among the known 57 genes annotated for senescence 35 are present within the 7190 DE genes (Fold enrichment= 2.01, p-value=1.46e-6, Table 3). Furthermore, 184 out of 286 predicted cellular senescence genes are among the 7190 DE genes (Fold enrichment= 2.10, p-value=2.89e-32, Table 3). This demonstrates that among the pool of differentially enriched genes between proliferating and senescent cells, genes predicted by PROTGOAT to be associated with cellular senescence, as well as those with current annotations, are significantly enriched. These results indicate that predictions made by PROTGOAT do reflect changes in the underlying biological process of cellular senescence.

**Table 1.**
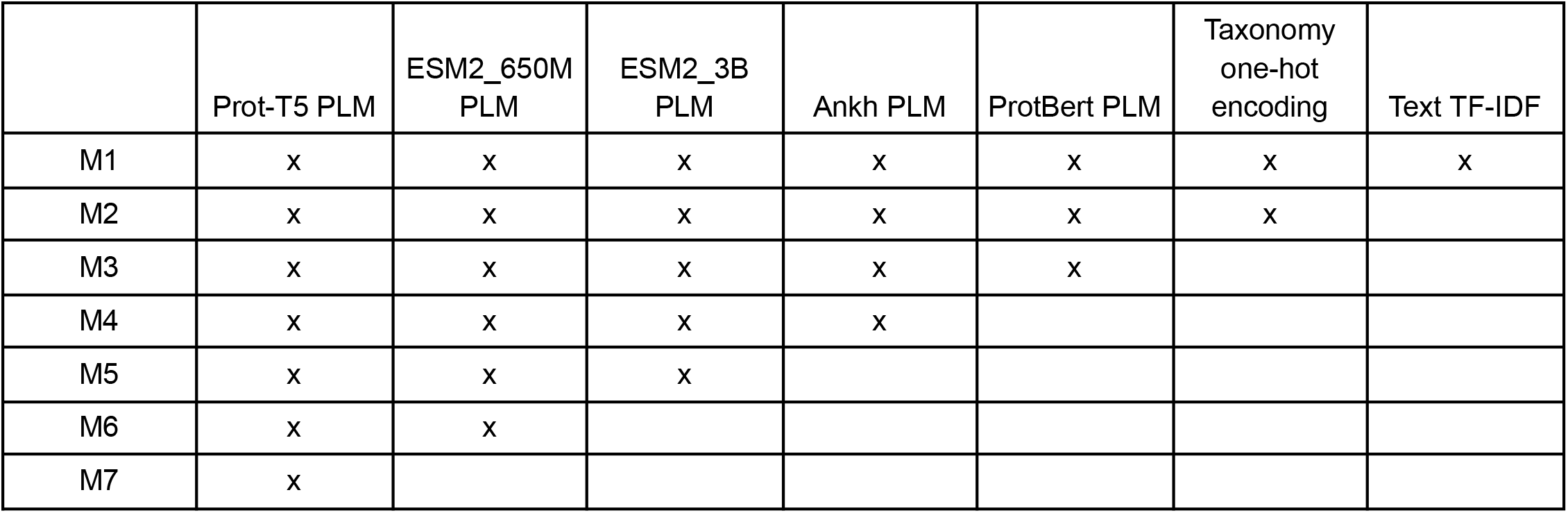
Components from PROTGOAT that were present within ablation study starting from the full model (M1)

|  | Prot-T5 PLM | ESM2_650M PLM | ESM2_3B PLM | Ankh PLM | ProtBert PLM | Taxonomy one-hot encoding | Text TF-IDF |
| --- | --- | --- | --- | --- | --- | --- | --- |
| M1 | x | x | x | x | x | x | x |
| M2 | x | x | x | x | x | x |  |
| M3 | x | x | x | x | x |  |  |
| M4 | x | x | x | x |  |  |  |
| M5 | x | x | x |  |  |  |  |
| M6 | x | x |  |  |  |  |  |
| M7 | x |  |  |  |  |  |  |

**Table 2.**
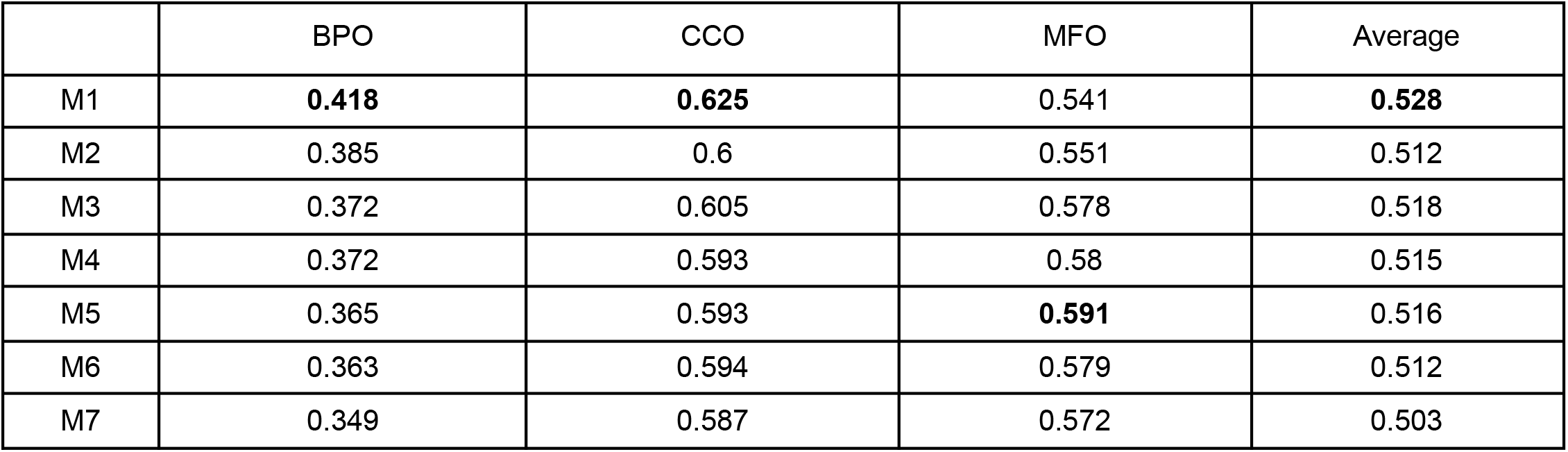
Change in F_max_ score for each GO domain upon progressive removal of different types of training data from PROTGOAT during training. Inference was carried out on the CAFA 5 test set.

|  | BPO | CCO | MFO | Average |
| --- | --- | --- | --- | --- |
| M1 | **0.418** | **0.625** | 0.541 | **0.528** |
| M2 | 0.385 | 0.6 | 0.551 | 0.512 |
| M3 | 0.372 | 0.605 | 0.578 | 0.518 |
| M4 | 0.372 | 0.593 | 0.58 | 0.515 |
| M5 | 0.365 | 0.593 | **0.591** | 0.516 |
| M6 | 0.363 | 0.594 | 0.579 | 0.512 |
| M7 | 0.349 | 0.587 | 0.572 | 0.503 |

**Table 3.** Calculated enrichment cores and p-values for genes predicted and currently annotated for cellular senescence as a proportion of differentially enriched genes from RNA-seq.

|  | Proportion of genes that are differentially expressed | Enrichment score | p-value |
| --- | --- | --- | --- |
| Annotated genes | 0.61 | 2.01 | 1.46e-6 |
| Predicted genes | 0.64 | 2.10 | 2.89e-32 |

## Discussion

We present here PROTGOAT, a new method for predicting the functions of proteins through Gene Ontology (GO) labels. We demonstrate through PROTGOAT that leveraging diverse pre-trained PLMs and integrating rich literature and taxonomy information leads to significant advancements in AFP. Notably, the success of PROTGOAT at the 5th Critical Assessment of Functional Annotation competition (CAFA 5), where it emerged as the 4th best model for predicting novel GO labels for proteins out of over 1600 submitted models, is indicative of the effectiveness of PROTGOAT against other methods for GO prediction.

The transformer-based PLMs at the core of PROTGOAT have harnessed the sequential nature of amino acids to draw powerful insights into protein functionality. Their ability to process and learn from extensive datasets that can contain billions of protein sequences has produced models that excel at recognizing nuanced patterns within protein sequences. Furthermore, by amalgamating this with curated textual information, PROTGOAT integrates the ‘literature knowledge’ of the scientific community into its prediction matrices.

We have demonstrated PROTGOAT’s competitive edge over existing state-of-the-art models based on their performance on the CAFA 5 test set. PROTGOAT’s ensemble approach mitigates the limitations of singular model dependency and captures a wider representation of protein functionalities. However, the ablation study reveals a domain-specific response to training data variance. The decline in performance within the BPO and CCO domain against the data ablation, and the unexpected improvement in the MFO domain, invites further investigation into the interaction between model complexity and domain specificity. These findings suggest that while a comprehensive dataset bolsters prediction accuracy in general, there may be an intricate balance between data sufficiency and overfitting and further work is needed to prevent overfitting of the model.

Moreover, we validate PROTGOAT’s performance and underscore its utility through identifying novel senescent related genes and proteins that are enriched in RNA-seq datasets of senescent cells. This highlights how the use of AFP tools like PROTGOAT allows for the rapid identification of candidate components and regulators of diverse biological processes, allowing for rapid hypothesis generation and target selection within experimental settings. However, it is also crucial to recognize that the enrichment of predicted genes in differentially expressed gene sets, while suggestive of relevance, does not equate to a direct functional role. Differential gene expression can be indicative of a protein’s involvement in a process, but it does not confirm a causal relationship. The genes predicted by PROTGOAT to be involved in senescence, while enriched, may include both drivers and passengers of this process. As such, further investigation is necessary to elucidate the specific contributions of these genes to the senescence phenotype. Each predicted gene represents a hypothesis generated by PROTGOAT that now requires experimental scrutiny.

While PROTGOAT marks a significant milestone in improvements of AFP technique, the journey towards a complete understanding of protein functionality is far from over, especially given the many overlapping and context-specific functions that proteins carry out. Future directions could include exploring alternative deep learning architectures that might better capture the hierarchical and graph-structured nature of protein functions, such as Graph Neural Networks (GNNs). These networks could potentially leverage the inherent structure of GO’s DAGs to better understand both the explicit and implicit relational context between different GO terms. Improvements to the way GO terms are annotated can also be a way to better reflect the complex nature and functions of proteins, potentially including the use of context specific GO terms or integrating information on dependency relationships within biological pathways. The additional dimensions such information adds to the available public training datasets will go a long way towards improving the performance of future AFP models.

In summary, PROTGOAT represents a significant leap forward in the AFP landscape, and the insights gleaned from this work pave the way for a new generation of AFP tools. As the biological research community continues to accumulate sequence data at an unprecedented rate, the deployment of sophisticated tools like PROTGOAT is imperative for transforming this wealth of data into actionable biological understanding.

## Author contributions

ZMC developed the PROTGOAT model and carried out training, inference, ablation, comparisons, and submissions to CAFA 5. AR carried out functional validation of senescence. ZMC generated embeddings from PLMs and AR generated embeddings from text data. ZMC wrote the main manuscript text and together with AR prepared all figures and tables. SS and PA proofread the manuscript. All authors reviewed the manuscript. All authors read and approved the final manuscript.

## Code availability

The code that was used to generate PROTGOAT can be found in this public GitHub repository : https://github.com/zongmingchua/cafa5

## Data availability

The embeddings generated for all 142246 proteins in the training set and 141864 proteins in the test superset of CAFA 5 are available for download at https://www.kaggle.com/datasets/zmcxjt/cafa5-train-test-data

## Acknowledgements

Numerous public notebooks on Kaggle served as inspirations for the final PROTGOAT model and were important for understanding of the CAFA 5 problem, in particular those by Antonina Dolgorukova, MT, Alexander Chervov, and Sergei Fironov. We are grateful for the help of Michael Alcaraz and Marcos Garcia Teneche in coming up with the name “PROTGOAT”.

